# The Vap33/Eph/Vav/Cdc42 complex confers temporal specification to the outgrowth of primary dendrites in Drosophila neurons

**DOI:** 10.1101/2022.09.19.508514

**Authors:** Daichi Kamiyama, Yuri Nishida, Rie Kamiyama, Miyuki Fitch, Takahiro Chihara

## Abstract

**SUMMARY:** The formation of primary dendrites (dendritogenesis) significantly affects the overall orientation and coverage of dendritic arborization, limiting the number and types of inputs a neuron can receive. Previously we reported how a *Drosophila* motoneuron spatially controls the positioning of dendritogenesis through the Dscam1/Dock/Pak1 pathway; however, how the neuron defines the timing of this process remains elusive. Here we show that the Eph receptor tyrosine kinase provides a temporal cue. We find that, at the onset of dendritogenesis, the Eph receptor recruits the Rho Family GEF Vav to the intracellular domain of Eph, which transiently activates the Cdc42 family of small GTPase. We also show that *vap33* (vesicle-associated membrane protein-associated protein) mutants exhibit defects in Cdc42 activation and dendritic outgrowth, indicating Vap33 may play an upstream role in Eph signaling. Together, our result and previous studies argue that the formation of primary dendrites requires the proximity of active Cdc42 and membrane-anchored Pak1 driven by collaborative action between two distinct signaling complexes, Vap33/Eph/Vav and Dscam1/Dock. Signal integration from multiple input pathways would represent a general mechanism for the spatiotemporal precision of dendrite branch formation.

## INTRODUCTION

The overall orientation and coverage of dendrites limit the number and types of inputs that a neuron can receive (Jan and Jan, 2003; 2010; Whitford et al., 2002). The pattern of dendritic arbors determines the integration of synaptic signals to the cell body and the firing pattern of an output signal (Jan and Jan, 2001). It is thus critical to understand how dendrites acquire their unique position and orientation in development. A mature dendritic morphology is acquired through multiple consecutive steps: In the initial step, primary dendrites extend away from the cell body to target fields (Kim and Chiba, 2004; Whitford *et al*., 2002); In the elongation step, dendrites undergo branching, extension, and retraction events to cover the target fields (Jan and Jan, 2010; Tavosanis, 2021); In the final step, self-avoidance and tiling mechanisms prevent dendritic receptive fields from overlapping with neighbors (Grueber and Sagasti, 2010; Parrish et al., 2007; Zipursky and Grueber, 2013). Terminal and intermediate branches are patterned by a combination of genetic instructions, substrate interactions, and neuronal activity (Williams et al., 2010; Wong and Ghosh, 2002). Over the past decades, extracellular signals, cell surface receptors, and intracellular regulators were identified to size and shape higher-order branches across different systems (Dong et al., 2015). In contrast, we have only rudimentary knowledge about the biological mechanisms of the first step of the morphological development of dendrites (i.e., dendritogenesis).

Previous studies using hippocampal neuronal cultures demonstrated how the outgrowth of primary dendrites is induced through intracellular genetic programs (Barnes and Polleux, 2009; Cheng and Poo, 2012; Craig and Banker, 1994). However, the cultures might not fully recapitulate developmental processes in the brain. Although current *in vivo* research in this area is limited, there is some evidence that extracellular signals influence dendritogenesis (Dong *et al*., 2015; Lin et al., 2020). For example, recent studies on *C. elegans* suggest that the Wnt signaling pathway initiates the formation of primary dendrites in an oxygen sensory neuron (PQR) (Kirszenblat et al., 2011). The Wnt ligand, LIN-44, is expressed from the tip of the animal tail. This ligand becomes an attractive signal to direct PQR dendrite outgrowth through the LIN-17/Frizzled receptor. In vertebrates, the class 3 secreted Semaphorin-3A (Sema3A) and its receptors neuropilin-1/plexin A promote the outgrowth of primary dendrites in hippocampal neurons (Polleux et al., 2000). The response of primary dendrites to Sema3A results in asymmetric localization of soluble guanylate cyclase to dendrites (Shelly et al., 2010). As a result, a hippocampal neuron extends primary dendrites from the cell body toward the pial surface. In *Drosophila*, we have previously performed a candidate-based screen for guidance receptors that affect dendrite morphogenesis (Furrer et al., 2007; Kamiyama et al., 2015). We found that while *Drosophila* embryos with a deletion in either the *frizzled* or *plexin A* gene show severe axon guidance defects, the same neurons grow dendrites normally. These results strongly suggest that other receptors promote the outgrowth of primary dendrites in *Drosophila*.

A set of 36 *Drosophila* embryonic motoneurons are present in each hemi-segment of the ventral nerve cord (Keshishian et al., 1996). Each of these neurons projects characteristic dendritic arbors and extends a single axon; additionally, individual motor axons innervate peripheral target muscles (**Figure 1A**). These motoneurons can be used to study every step of the morphological changes during development. For example, the aCC (anterior corner cell) motoneuron has been used to study axon growth and guidance, dendrite elaboration, and synapse formation (Ackerman et al., 2021; Furrer et al., 2003; Fushiki et al., 2016; Schneider-Mizell et al., 2016; Tripodi et al., 2008; Wolf and Chiba, 2000). In particular, because this neuron sends out the primary dendrites at a stereotypical position on the axon shaft (**Figure 1B**), it is well-suited for investigating the molecular mechanisms of the precise branching pattern of primary dendrites (Kamiyama and Chiba, 2009; Kamiyama *et al*., 2015). Our genetic and imaging analyses have previously demonstrated that, upon contact with the MP1 pioneer neuron, the aCC motoneuron exhibits initial accumulation of the Down syndrome adhesion molecule (Dscam1) at the contact site from 11 hours after egg laying (11:00 AEL) (**Figure S1A top**) (Kamiyama *et al*., 2015). This interaction sets up the site of dendritic outgrowth in aCC. Accumulated Dscam1 consequently acts via the Dreadlocks (Dock) adaptor protein and recruits the p21-activated kinase (Pak1) to the plasma membrane. Within the next two hours (13:00 AEL), the Dscam1/Dock/Pak1 complex initiates the outgrowth of primary dendrites via the small GTPase Cdc42 (**Figure S1A** and **S1B**).

**Figure 1.**
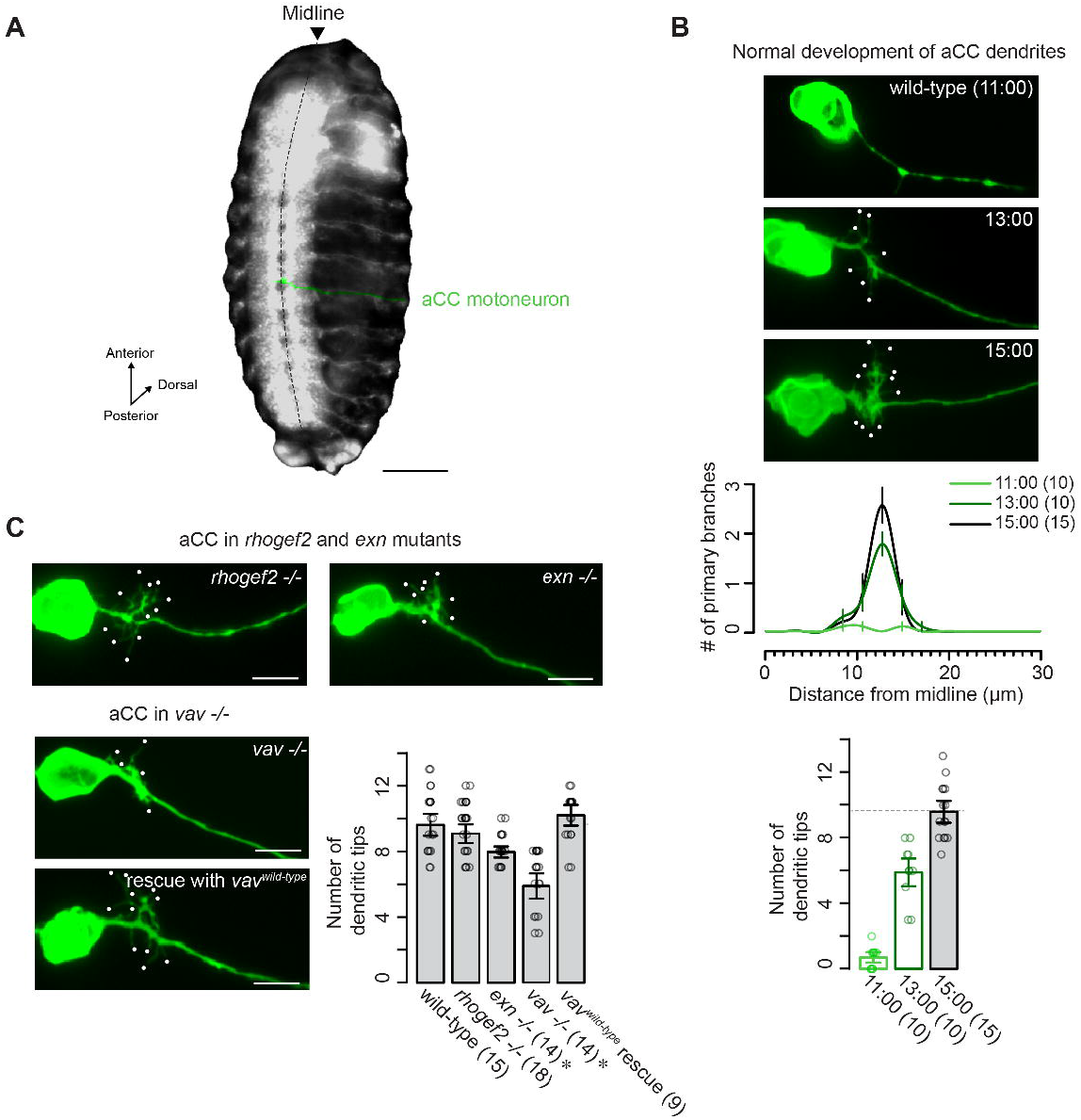
The *vav* gene is required for the outgrowth of primary dendrites in the aCC motoneuron. (**A**) Representative image showing the location of the aCC motoneuron (green) in an embryo at 15:00. Ventral view with anterior up. (**B**) Top: Normal development of aCC in wild-type embryos. Developmental stages (11:00, 13:00, and 15:00 AEL) are indicated in each panel. Anterior is up, and dorsal is to the right in these and subsequent images. Middle and bottom: Quantifications of the distribution of primary tips and the number of dendritic tips. Sample sizes are indicated in parentheses. Data are represented as mean ± SEM. (**C**) Top and left: Representative images of dendritic outgrowth in *rhogef2* null, *exn* null, *vav* null, and *vav* wild-type rescue embryos at 15:00. Right: Quantification of the number of dendritic tips. A two-tailed Student’s t-test between a control sample (wild-type) and each mutant sample was used for statistical analysis (^*^, *p* < 0.05). Scale bars, 100µm in (**A**) and 5µm in (**C**).

The loss of the *cdc42* gene leads to a drastic reduction of dendritic tips in aCC; therefore, Cdc42 is a crucial regulator of aCC dendritogenesis (Kamiyama and Chiba, 2009). Cdc42 acts as a molecular switch between an active GTP-bound form and an inactive GDP-bound form, regulated by guanine nucleotide exchange factors (GEFs) and GTPase activating proteins (GAPs), respectively (Hall, 1998). These GEFs and GAPs are reciprocally regulated by upstream signaling molecules. When bound to GTP, Cdc42 binds to downstream effectors, including Pak1, that transduce signals to control the actin cytoskeleton (Bishop and Hall, 2000). Knowing the role of Cdc42 in linking its activation to downstream effectors, we have previously investigated the pattern of Cdc42 activation in aCC. We have engineered a FRET-based activation probe for Cdc42 (Cdc42 <u>a</u>ctivation <u>probe</u>, also called a-Probe) (Kamiyama and Chiba, 2009), which uses Cdc42 tethered by a flexible linker to the Cdc42-binding domain (CBD) of Pak1 (**Figure S1C**). Because Cdc42 activation increases the intramolecular binding between Cdc42 and CBD, the resulting conformation changes induce FRET between a flanking CFP and YFP pair. Imaging studies with this probe have revealed that the time course of Cdc42 activation correlates with dendrite development. We have observed that sustained Cdc42 activity near the site of primary dendrites is induced from 13:00 AEL. The spatial proximity of GTP-bound Cdc42 and membrane-anchored Pak1 would allow the immediate activation of other cytoskeletal regulatory proteins, promoting efficient actin polymerization. The aCC motoneuron then could undergo a process that projects dendritic branches from the axon shaft at 13:00 AEL (**Figure S1A** and **S1B**). Although it has been reported that Cdc42 is regulated in response to environmental cues, little is known about the major components of an upstream signal mediating the activation of Cdc42 to initiate dendritic outgrowth.

In this paper, we report the characterization of Vav, the Dbl family of GEF, which acts as an upstream component of Cdc42. Vav is recruited by the Eph receptor tyrosine kinase to the plasma membrane, and this translocation event is critical to catalyzing the exchange of GDP for GTP on Cdc42. Moreover, we identify a putative ligand for dendritic outgrowth mediated by the Eph receptor and demonstrate that a deletion mutant of the ligand, Vap33, largely suppresses the Cdc42-dependent outgrowth of primary dendrites. We envision that upon ligand binding, Eph undergoes dimerization and tyrosine autophosphorylation. Vav then associates with Eph and activates Cdc42 on the membrane. Together, these data reveal how two distinct signaling pathways, Vap33/Eph/Vav and Dscam1/Dock, coordinate Cdc42 activity and Pak1 localization to drive the formation of primary dendrites in a spatiotemporal manner.

## RESULTS

### Abnormal aCC dendritogenesis in the *vav mutant* embryos

In previous studies, we have quantitatively characterized dendrite morphogenesis in the aCC motoneuron (Kamiyama and Chiba, 2009; Kamiyama *et al*., 2015). We used a method that allowed retrograde labeling of aCC (Inal et al., 2020) and imaged development from 11 to 15 hours after egg laying (AEL). We then quantified the number of dendritic tips and the position of primary dendritic tips (**Figure 1B**). We found that the initial outgrowth of the aCC dendritic arbor starts at 13 µm from the midline at 13:00 AEL (**Figure 1B**). This process is highly stereotyped between the aCC motoneurons in different embryos. We also demonstrated that mutations in *cdc42* cause a significant decrease in the number of aCC dendritic tips after 13:00 AEL, while no defects were observed in aCC before 13:00 AEL (Kamiyama and Chiba, 2009). In addition, the phenotype was rescued by an aCC-specific expression of *cdc42* (Kamiyama and Chiba, 2009). These data suggested a cell-autonomous role of *cdc42* in aCC dendritogenesis. It is interesting to note that *rac* loss-of-function mutations (*rac1, rac2*, and *mig-2-like*), which are closely related to *cdc42*, exhibited no gross morphological defects at these developmental time points (Kamiyama *et al*., 2015). To further elucidate the mechanisms by which Cdc42 promotes dendritic outgrowth, we have previously engineered a FRET-based activation probe for Cdc42, also called aProbe (Kamiyama and Chiba, 2009). Measuring the pattern of Cdc42 activation with aProbe, we found that it is restricted to the site of dendritic outgrowth at the onset time of dendritogenesis (see also **Figure 2A**). These findings raised the possibility that there might be molecules that play an upstream role in spatiotemporally activating Cdc42. For example, in regulating cell polarity in yeast, the restricted localization of upstream components (i.e., GEFs) is critical to confine Cdc42 activation (Chiou et al., 2017).

**Figure 2.**
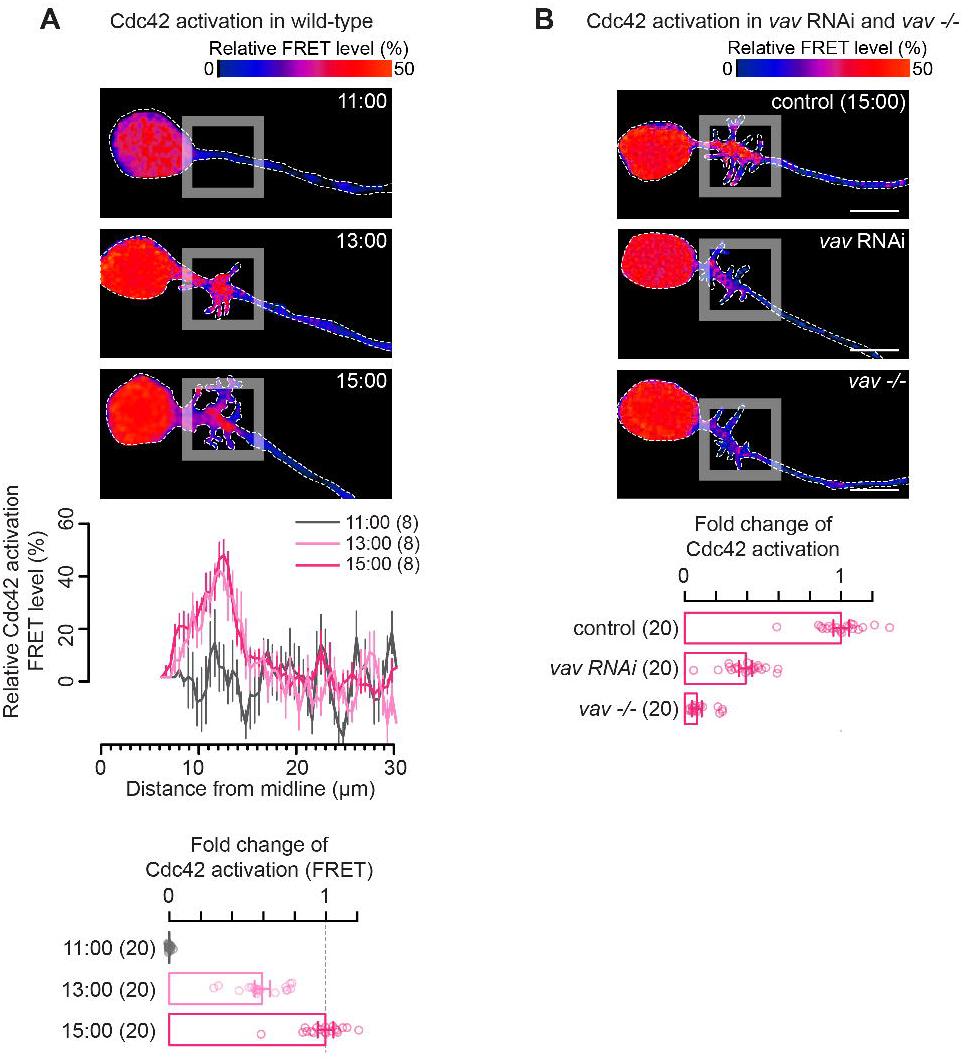
Cdc42 is activated by Vav in aCC. (**A**) Top: Developmental time course of aCC expressing a *UAS-aProbe* construct driven by *eve-GAL4*. Pseudocolor images show measured FRET signals. Warmer colors correspond to higher FRET signals. White dashed lines outline individual neurons. Middle: Traces show the spatiotemporal changes of Cdc42 activity (each trace is a combined average of eight neurons). Bottom: Quantification of Cdc42 activation at different developmental time points, showing the fold change in FRET levels. (**B**) Top: Pseudocolor images of aProbe patterns in control, *vav* RNAi, and *vav* null embryos at 15:00. Bottom: Quantification of Cdc42 activation in a region where aCC dendrites grow. Data are shown as mean ± SEM. Scale bars, 5µm.

As a step toward understanding Cdc42 activation mechanisms that regulate aCC dendritogenesis, we began with screening of RNA interference (RNAi) constructs for all genes in the *Drosophila* genome encoding Cdc42 GEFs (**Figure S1D**). *UAS-RNAi* constructs were expressed in aCC and its sibling RP2 and pCC neurons using a specific *GAL4* line (an *even-skipped-GAL4* coding fusion transgene). The *GAL4* transgene was expressed 4 hours prior to dendritogenesis and continued throughout embryogenesis (Kamiyama *et al*., 2015). Phenotypes in aCC dendritogenesis were determined at 15:00 AEL. Knocking down any one of the genes encoding Ephexin (Exn), Vav, and RhoGEF2 exhibited a significant reduction in the dendrite tip number, which is similar to the *cdc42* RNAi case (**Figure S1D**). To further validate their role in aCC, we imaged aCC dendritogenesis in embryos homozygous for loss-of-function mutations in *exn, vav*, or *rhogef2*. Because genetic deletion of *rhogef2* did not affect aCC dendritogenesis at all (**Figure 1C**), the RNAi-induced defect is likely due to the knockdown of off-target genes. Between the two GEFs (*exn* and *vav*), the *vav* knockout caused the most severe reduction in the dendrite tip number (**Figure 1C**). While suspecting functional redundancy between the two, we here focused on *vav*, which encodes a cytosolic protein that belongs to the Dbl GEF superfamily. In contrast to vertebrates that have multiple *vavs* (*vav1, vav2*, and *vav3*), *Drosophila* has a single *vav* (Rodriguez-Fdez and Bustelo, 2019). Previously, it was shown that *Drosophila vav* is necessary in a subset of interneurons for axonal growth within the embryonic CNS (Malartre et al., 2010). Contrary to those interneurons, *vav* mutation does not influence aCC motor axon formation but leads to defects specifically in dendrite morphogenesis. This aCC loss-of-dendrite phenotype in *vav* null mutants was rescued by selectively resupplying a *vav* cDNA in aCC (**Figure 1C**). Taken together, the aCC-specific knockdown and rescue experiments indicate that Vav acts within aCC to initiate dendritic outgrowth in a cell-autonomous manner.

### Cdc42 GTPase is activated by endogenous Vav in the aCC motoneurons

As shown in a different animal model (Cowan et al., 2005), Vav appears to be a key regulator of Rac activation in axon guidance (see also the **DISCUSSION** section). Because Rac and Cdc42 often share upstream regulators, it is conceivable that Vav may be employed for Cdc42 to regulate other aspects of neuronal development. To test this possibility, we assessed the effect of *vav* knockdown on the activation of Cdc42 in aCC. To visualize Cdc42 activation with single-cell resolution, we used transgenic flies that express aProbe under the control of the *GAL4/UAS* system. When expressed by *eve-GAL4*, aProbe exhibited a FRET signal enriched in the proximal region of the aCC axon. A quantitative analysis of the FRET signal in different time points (i.e., 11:00, 13:00, and 15:00 AEL) showed that Cdc42 starts to be active at 13:00 AEL, consistent with the onset time point for dendritic outgrowth (**Figure 2A**). To assess whether Vav is required for Cdc42 activation at the region, we measured FRET signals in both control and *vav* RNAi-expressing aCC from 13:00 to 15:00 AEL. In aCC expressing *UAS-vav RNAi*, Cdc42 activation is significantly reduced during our observation period (**Figure 2B**). However, the penetrance of the *vav* RNAi phenotype was relatively low (< 60%) and might be due to an incomplete knockdown of *vav* in aCC, as aCC displayed a much more substantial reduction in Cdc42 activity in null mutants (**Figure 2B**). These findings demonstrate that Vav is the primary GEFs to regulate the activation of Cdc42 during aCC dendritogenesis.

### Vav is expressed differentially in the aCC motoneurons

Previous RNA *in situ* hybridization studies showed that *vav* appeared to be ubiquitously expressed throughout most tissues in the *Drosophila* embryo (Malartre *et al*., 2010). To visualize the distribution of Vav proteins, we modified the endogenous *vav* genomic locus in a BAC (bacterial artificial chromosome) via a recombineering approach (see **EXPERIMENTAL PROCEDURES**) and inserted a GFP tag upstream of the *vav* coding sequence (*GFP-vav*; see also **Figure S2A**). The modified BAC was introduced into flies as a transgene (**Figure S2B**). Because the BAC partially rescues lethality in *vav* mutants (data not shown), the BAC-encoded GFP-Vav is likely to function similarly to the unmodified, endogenous protein. Our imaging analysis shows that, although tagged Vav displays an even distribution in early embryonic stages, it starts to accumulate in the longitudinal connectives from the onset of dendritic outgrowth (i.e., 13:00 AEL) and remains detectable until the end of embryogenesis (**Figure S2C**).

We next sought to visualize Vav localization specifically within the aCC motoneurons. However, it is challenging because Vav is broadly present in cellular processes that intertwine with aCC dendrites. As failing to show aCC-specific localization of Vav, we imaged Vav-GFP in embryos that express a *UAS-vav-GFP* transgene using aCC-specific *eve-GAL4*. Because aCC-specific expression of Vav-GFP is sufficient to rescue defective dendritic outgrowth in *vav* mutants (**Figure 1C**), the distribution of overexpressed Vav-GFP would potentially reflect the endogenous localization of Vav during aCC dendritogenesis. Under this condition, we found that Vav-GFP is enriched at the region at 13:00 AEL when Cdc42 is usually activated (**Figure 3**). Importantly, such a confined localization of Vav started at the onset of dendritic outgrowth, spatiotemporally coinciding with the Cdc42 activation pattern.

**Figure 3.**
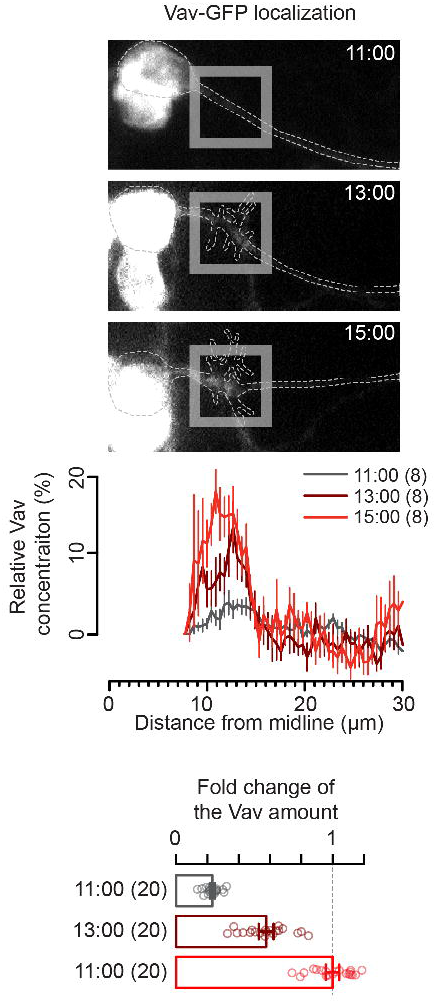
Vav is enriched at the site of aCC dendritogenesis. Top: Representative images of developing aCC in animals that express wild-type *vav-GFP*. Middle: Traces of Vav-GFP distribution. Bottom: Quantification of Vav enrichment before and after dendrite initiation. GFP intensity was measured at the site of aCC dendritogenesis. Data are shown as mean ± SEM.

### The GEF Vav interacts with the intracellular domain of Eph in *Drosophila* S2 cells

Earlier biochemical studies provided a basis for a model of *vav* function in the mammalian brain during the period when axon guidance happens (Cowan *et al*., 2005). Recruitment of Vav2 to the plasma membrane in response to Eph receptor tyrosine kinase activation could promote the exchange of bound GDP for GTP and thus activate the Rac family of small GTPase (Cowan *et al*., 2005). This translocation happens through direct binding of Vav2 to the EphA4 receptor (Cowan *et al*., 2005). An interaction site has been mapped to the juxtamembrane (JM) tyrosine residues in EphA4 and the SH2 domain of Vav2 (Cowan *et al*., 2005). In contrast to mammals, the *Drosophila* genome encodes a single Eph receptor (Morrison et al., 2000). *Drosophila* Eph includes four tyrosine residues in the JM region related to EphA4 (**Figure S3**). Vav was tested for its ability to associate with Eph. We transiently transfected *Drosophila* S2 cells with expression plasmids encoding FLAG-tagged Vav (a FLAG-tag fused to Vav wild-type) and Eph-IC-Fc (a fusion protein containing the human Fc domain and intracellular domain of Eph) (**Figure 4A**). Then we immunoprecipitated Eph receptors with protein A beads. The immunoprecipitants were subsequently analyzed on Western blots. When overexpressed in cells, Eph-IC would be active presumably because of high expression levels (Cowan *et al*., 2005).

**Figure 4.**
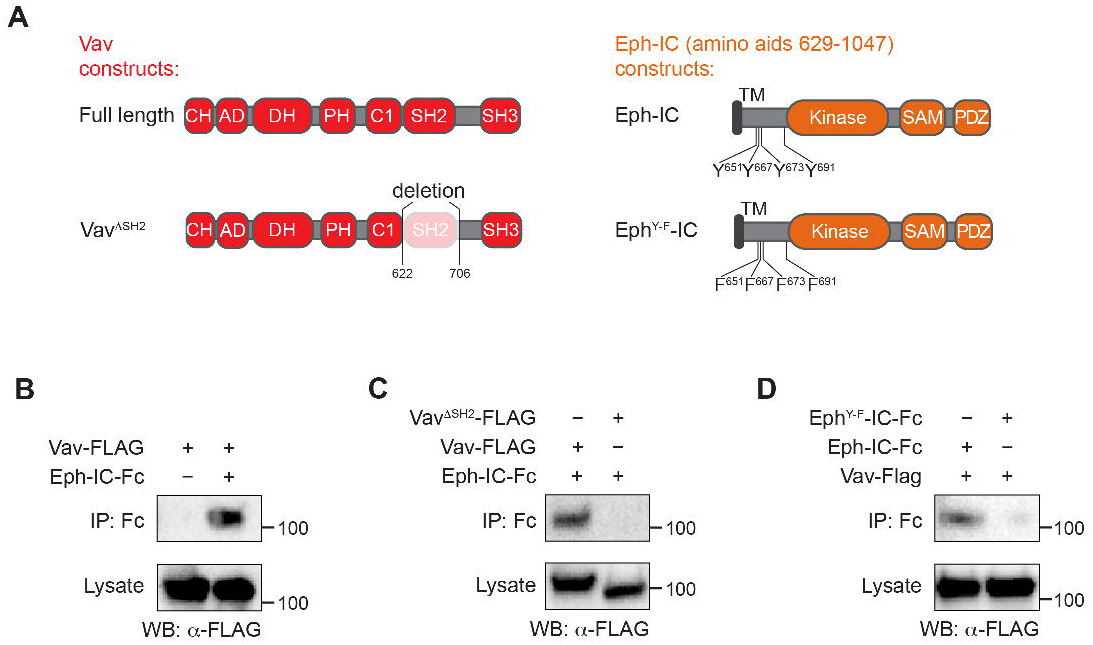
Vav physically interacts with the Eph receptor tyrosine kinase in *Drosophila* S2 cells. (**A**) Cartoon representation of Vav and Eph constructs used in Co-IP assays. The numbers refer to amino acid positions. TM: transmembrane domain. IC: intracellular domain. (**B**-**D**) Western blot images showing Co-IP between Vav-FLAG and Eph-Fc. IP: immunoprecipitation. WB: western blot. Gel images are representative of at least two independent repeats.

First, we confirmed that Vav and Eph-IC physically interact (**Figure 4B**). Second, we demonstrated that the SH2 domain of Vav is essential for interaction with Eph. Vav^ΔSH2^ (i.e., this is a Vav deletion construct lacking the SH2 domain) did not interact with Eph-IC (**Figure 4C**). Third, we demonstrated that Vav interacts with the JM tyrosines of Eph. Vav failed to interact with an Eph JM tyrosine mutant (Y651F/Y667F/Y673F/Y691F; **Figure 4D**). In summary, we observed an interaction between the JM tyrosines of Eph and the SH2 domain of Vav. These data are similar to the interaction observed previously in mammals (Cowan *et al*., 2005).

### The Eph receptor activates Cdc42 in the aCC motoneurons

As described above, Vav physically binds to the Eph receptor in *Drosophila* S2 cells. This receptor is known to play a critical role in a wide range of neuronal processes, including local dendrite branch dynamics, axon navigation, and synapse formation in *Drosophila* (Anzo et al., 2017; Boyle et al., 2006; Frank et al., 2009; Sardana et al., 2018). We thus sought to investigate whether Eph controls dendritic outgrowth in the aCC motoneurons. To this end, we counted the number of dendritic tips in *eph*^*x652*^ homozygous mutants (*eph*^*x652*^ appears to be a protein null (Boyle *et al*., 2006)). Homozygous *eph*^*x652*^ mutant embryos significantly reduced the aCC dendrite tip number indistinguishable from those observed in *vav* mutants (**Figure 5A**). The phenotype was rescued by aCC-specific expression of a wild-type *eph* cDNA under the control of *eve-GAL4* (**Figure 5A**). Likewise, aCC-specific *eph* RNAi caused a severe reduction in the dendrite tip number, phenocopying an *eph* null mutation (**Figure 5A**). These data indicate that Eph functions aCC dendritogenesis in a cell-autonomous manner. Having demonstrated the similarity of the *vav* and *eph* mutant phenotype, we set out to assess whether Eph and Vav act within the same signaling pathway. To explore a functional interplay between *vav* and *eph*, we generated embryos heterozygous for both *vav* and *eph*. We found that the doubly heterozygous mutations caused a severe impairment (32% of reduction of the dendrite tip number) of dendritic outgrowth (**Figure S4A**). In contrast, *vav +/-* or *eph +/-* heterozygous mutation did not affect dendritogenesis (**Figure S4A**). Such a synergistic genetic interaction between heterozygous mutations can be taken as evidence that they act together to initiate dendritic outgrowth.

**Figure 5.**
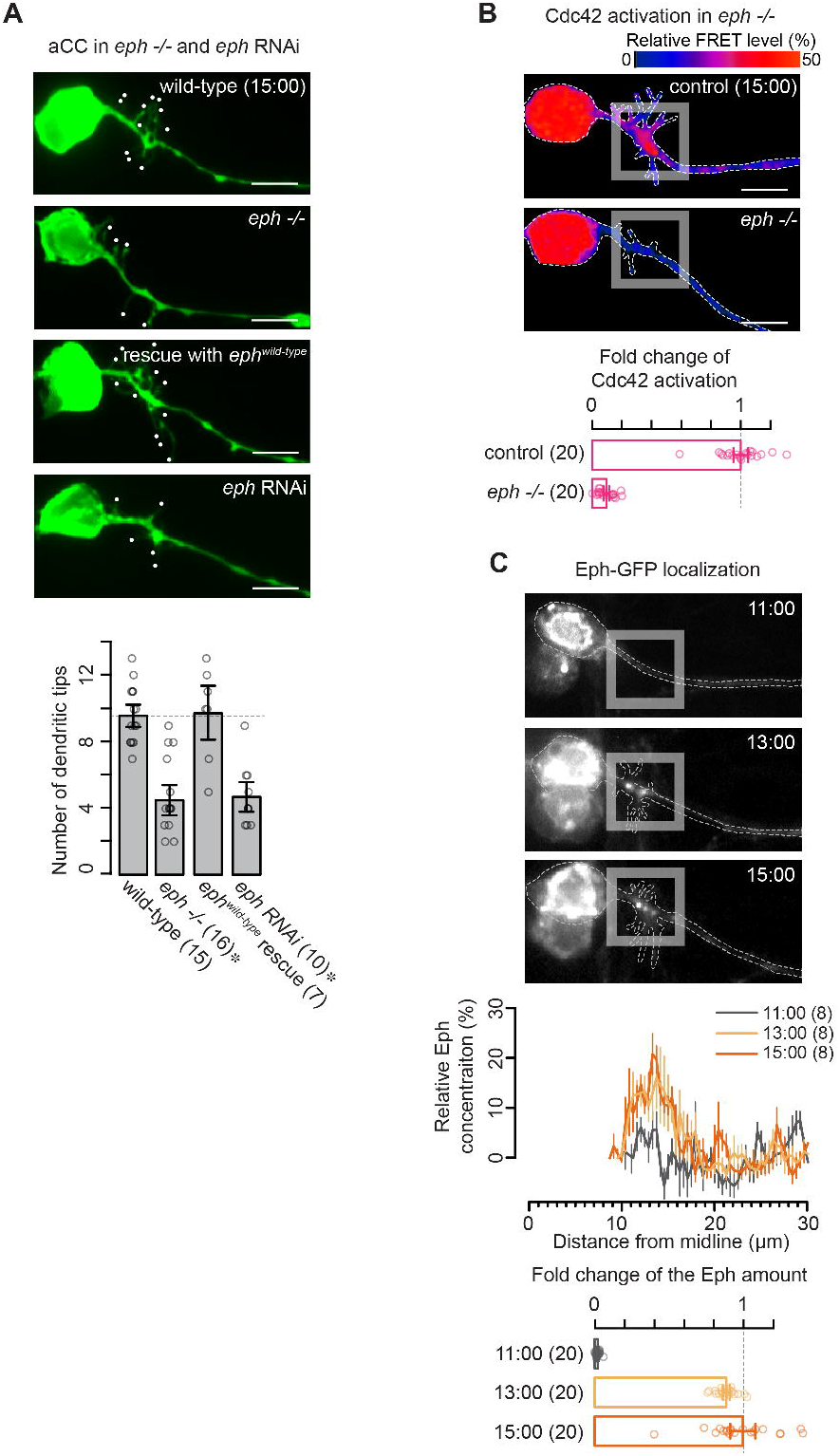
Eph is required for the outgrowth of primary dendrites. (**A**) Top: Representative images of dendritic outgrowth in wild-type, *eph* null, *eph* wild-type rescue, and *eph* RNAi embryos at 15:00. Bottom: Quantification of the number of dendritic tips. Statistical comparison was carried out to a control sample (wild-type) with a two-tailed Student’s t-test (^*^, *p* < 0.05). (**B**) Top: Pseudocolor images of aProbe responses in control and *eph* null embryos. Bottom: Quantification of the level of Cdc42 activation per aCC for the indicated genotypes. (**C**) Top: Representative images of aCC expressing an Eph-GFP fusion driven by *eve-GAL4* in different time points. Middle: Traces of Eph-GFP distribution. Bottom: Quantification of Eph enrichment before and after dendrite initiation. Data are shown as mean ± SEM. Scale bars, 5µm in (**A**) and (**B**).

To gain insight into the role of Eph in aCC dendritogenesis, we examined the pattern of Cdc42 activation in *eph*^*x652*^ mutants using the aProbe reporter. If the Eph receptor could function together with Vav, disrupting *eph* would alter the level and spatial extent of Cdc42 activation in these neurons. Our observation confirms this prediction; Cdc42 activation during aCC dendritogenesis is almost abolished in *eph*^*x652*^ mutants (**Figure 5B**). Thus, Eph receptor regulation, together with Vav, plays an essential role in modulating Cdc42 activity to orchestrate the outgrowth of primary dendrites.

### Eph receptors are enriched at the site of dendritic outgrowth and developmentally regulated

To this point, our genetic and biochemical data are compatible with a model in which Eph acts with Vav to activate Cdc42 at the proximal region of the axon when aCC dendrites start to emerge. However, the data have not yet addressed when and where the *eph* gene is expressed. To determine if *eph* is expressed in the embryonic CNS at an appropriate time to play a role in dendritic outgrowth, we visualized the pattern of *eph* expression. We previously introduced a *myc* coding sequence into the *eph* gene locus by CRISPR/Cas9-mediated homology-directed repair (Anzo *et al*., 2017). This allele allows the visualization of endogenous Eph proteins *in vivo*. Incubation of fixed, dissected embryos with an anti-myc antibody reveals that *eph* is expressed in most CNS neurons at 13:00 AEL; Eph is detectable predominantly along the longitudinal connectives where aCC dendrites normally grow (**Figure S4B**).

Next, we sought to determine the subcellular localization of Eph in the aCC motoneuron. To this end, we expressed GFP-tagged Eph in these neurons. We examined Eph localization at different time points (i.e., 11:00, 13:00, and 15:00 AEL) (**Figure 5C**). Expression of Eph-GFP showed that it starts to localize at the site of dendritogenesis at 13:00 AEL when Cdc42 becomes activate. Note that the receptors form clusters at the region, indicating that the receptors are most likely active (Craig and Banker, 1994). The width of Eph distribution matches that of Vav distribution, consistent with the width of the region in which Cdc42 is usually activated (6.0 ± 1.2µm, 5.3 ± 0.6µm, and 5.9 ± 1.2µm at 13:00, respectively). These findings indicate that both Eph and Vav are present at an appropriate time and place that allows them to control Cdc42 activity in aCC.

### Characterization of the Eph/Vav interaction when promoting dendritic outgrowth

To further characterize the nature of the interaction between Vav and Eph, we sought to determine the domains of each protein required for aCC dendritogenesis. We generated a series of *GFP*-tagged deletion and mutation constructs and tested the ability to rescue the aCC loss-of-dendrite phenotype under the control of *eve-GAL4*. We confirmed beforehand that these constructs exhibited somatodendritic localization.

We first examined the requirement for the SH2 domain of Vav, which mediates its interaction with Eph in *Drosophila* S2 cells (see also **Figure 4C**). A Vav mutant lacking the SH2 domain failed to rescue defects in dendritic outgrowth (**Figure 6C, C’**, and **G**), indicating that the Vav SH2 domain is required for aCC dendritogenesis. How then can the Vav SH2 interact with Eph? Our biochemical studies indicate in S2 cells that the SH2 domain potentially binds to tyrosine residues phosphorylated on Eph. To assess the significance of Eph phosphorylation, we employed non-phosphorylatable mutations of tyrosine residues within the JM region of Eph. As the mutations abolished the Vav-Eph interaction in the co-IP assay (see also **Figure 4D**), the same mutant form did not rescue the aCC loss-of-dendrite phenotype in *eph* mutants (**Figure 6E, E’**, and **G**). These data suggest that those tyrosine residues in the JM region would be phosphorylated to promote dendritic outgrowth.

**Figure 6.**
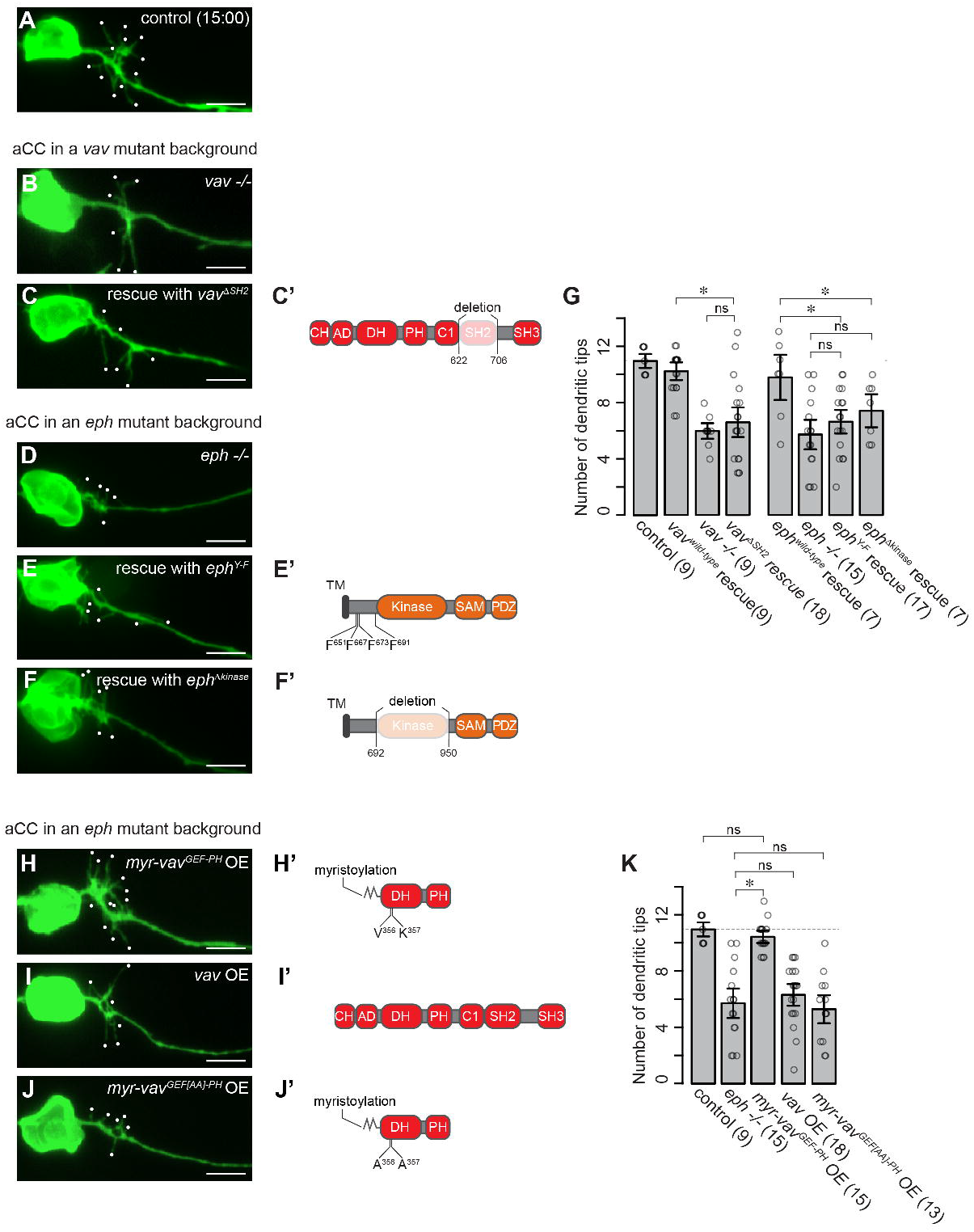
Interaction between Vav and Eph is required for dendritic outgrowth. (**A** - **F** and **H** - **J**) Representative images of dendritic outgrowth in control, *vav* null, *vav*^*ΔSH2*^ rescue, *eph* null, *eph*^*Y-F*^ rescue, and *eph*^*Δkinase*^ rescue, *myr-vav*^*GEF-PH*^ overexpression, *vav* wild-type overexpression, and *myr-vav*^*GEF[AA]-PH*^ overexpression embryos at 15:00. OE: overexpression. (**C’, E’, F’, H’, I’**, and **J’**) Schematics showing Vav and Eph constructs used in those experiments. (**G** and **K**) Quantification of the number of dendritic tips per aCC for the indicated genotypes. A one-way ANOVA was used for multiple comparisons (^*^, *p* < 0.05; ns, *p* > 0.05). Data are shown as mean ± SEM. Data are shown as mean ± SEM. Scale bars, 5µm.

The intracellular domain of Eph contains a tyrosine kinase domain, which presumably leads to Eph autophosphorylation on tyrosine residues in the JM region (Cowan *et al*., 2005). To determine whether the kinase domain is essential for aCC dendritogenesis, a deletion mutant was tested for the ability to rescue the *eph* dendritogenesis defects. A mutant form of Eph lacking the kinase domain failed to restore dendritic outgrowth in aCC (**Figure 6F, F’**, and **G**), indicating that the intrinsic kinase activity is necessary for aCC dendritogenesis. In summary, these data support a model in which Eph activation results in autophosphorylation of the JM tyrosines, generating docking sites for the SH2 domain of Vav.

### Vav membrane localization is sufficient to promote dendritic outgrowth

To seek additional evidence to support a role for Vav acting downstream from Eph, we generated a membrane-anchored form of Vav. A myristylation signal was fused to a catalytic RhoGEF DH-PH domain (myr-Vav^DH-PH^). If the major function of Eph is to anchor cytoplasmic Vav to the plasma membrane, then the expression of *myr-vav*^*DH-PH*^ might restore the *eph* mutant phenotype. To test this, we expressed *UAS-myr-vav*^*DH-PH*^ under the control of *eve-GAL4* in *eph*^*x652*^ homozygous mutants. This time, we overexpressed *myr-vav*^*DH-PH*^ at high enough levels to interfere with the function of the endogenous *vav* gene. To achieve the highest levels of its expression, we used embryos carrying two copies of *eve-GAL4* to drive the *UAS* transgene in aCC. In *eph* mutants, the dendrite branch number was severely reduced; in contrast, the *eph* phenotype was fully rescued by overexpression of *myr-vav*^*DH-PH*^ (**Figure 6H, H’**, and **K**). To exclude the possibility that the restoration of the number of dendrite tips is simply due to a higher amount of Vav, we assessed the ability of wild-type *vav* to rescue the *eph* phenotype. aCC-specific expression of wild-type *vav* did not rescue the phenotype (**Figure 6I, I’**, and **K**).

The rescue also requires GEF catalytic activity. We introduced a double-point mutation (L356A, K357A) into the DH domain of Vav, which is known to eliminate GEF catalytic activity (Martin-Bermudo et al., 2015). Importantly, high-level expression of this mutant form (*myr-vav*^*DH[A356A357]-PH*^) failed to rescue the *eph* phenotype (**Figure 6J, J’**, and **K**). These results argue that GEF activity on the plasma membrane is sufficient to promoting dendritic outgrowth.

### *vap33* is necessary for Eph-induced dendritic outgrowth of aCC

Many, if not all, Eph-dependent biological processes are triggered by its cognate ligands. Ephrin (Lisabeth et al., 2013; Taylor et al., 2017; Wilkinson, 2001) and Vap33 (vesicle-associated membrane protein-associated protein) (Tsuda et al., 2008) are the only known ligands for the Eph receptor. In the *Drosophila* brain, Eph mediates axon repulsion, synaptic homeostasis, and olfactory map formation through binding to Ephrin (Anzo *et al*., 2017; Boyle *et al*., 2006; Frank *et al*., 2009; Sardana *et al*., 2018) and axon guidance of the mushroom body neurons through binding to Vap33 (Tsuda *et al*., 2008). To assess the requirement of *ephrin* and *vap33* for dendritic outgrowth, we measured the number of aCC dendritic processes in *ephrin*^*I95*^ and *vap33*^*Δ31*^ mutant embryos. These two alleles are reported to be protein nulls (Anzo *et al*., 2017; Tsuda *et al*., 2008). We found that the dendrite tip number was significantly reduced in *vap33* mutants, but there was no significant difference between *ephrin* and wild-type animals (**Figure S5A**). Since *vap33* homozygous mutants show an obvious phenotype, we focus on the role of *vap33* for the rest of this study.

### The role for *vap33* in Eph-dependent aCC dendritogenesis

Previous work demonstrated that Vap33 is present in *Drosophila* larval tissues (Tsuda *et al*., 2008). To assess whether Vap33 could act as a ligand to promote dendritic outgrowth in aCC, we examined its distribution more precisely at embryonic stages. We used previously generated antibodies against Vap33, of which specificity has been thoroughly confirmed (Tsuda *et al*., 2008). Vap33 is expressed in the embryonic CNS as early as 10:00 AEL (data not shown).

Vap33 distribution within neurons was assessed in preparations double-stained with anti-Vap33 and a neuron marker (anti-horseradish peroxidase). Vap33 staining was observed in specific organelles, possibly the ER. In addition, staining was observed at cell-cell boundaries, indicating that the proteins are potentially in the extracellular space (data not shown). To examine the presence of extracellular Vap33, we used the optimized procedure for immunohistochemistry only to visualize extracellular Vap33 (Tsuda *et al*., 2008) and quantified the patterns of extracellular Vap33 distribution at 13:00 and 15:00 AEL. Between 13:00 and 15:00 AEL, the extracellular Vap33 signal is significantly increased in the longitudinal connectives where dendrites normally emerge (**Figure 7A**; note that no specific signal is detected prior to 13:00 AEL nor in *vap33* null mutants).

**Figure 7.**
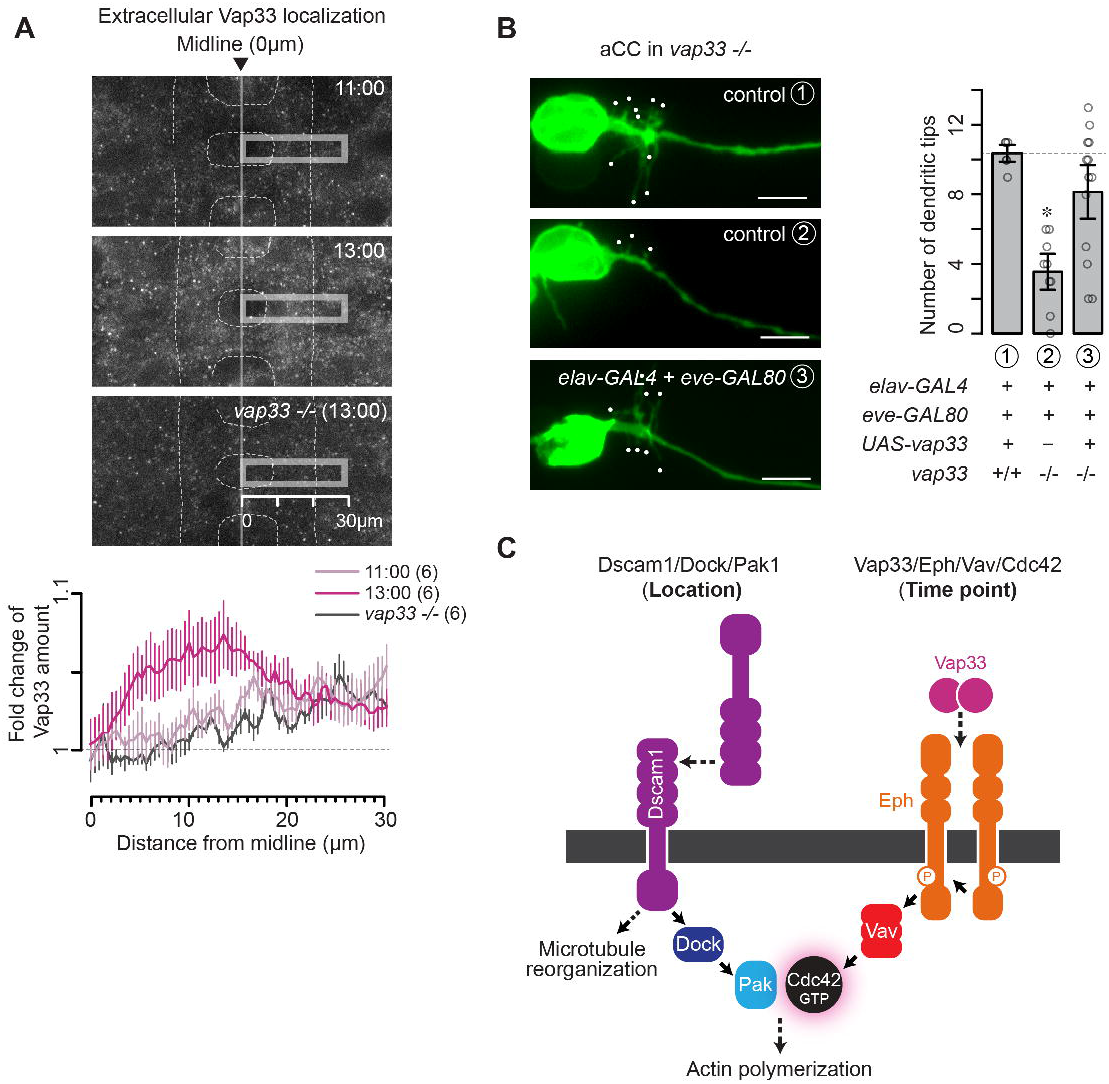
Vap33 functions non-cell-autonomously to control dendritic outgrowth in aCC. (**A**) Top: Fluorescence images of anti-Vap33 staining of wild-type embryos at the indicated developmental stages. The fluorescence signal is almost diminished in *vap33* mutants. Neuropiles are outlined by white dash lines. Bottom: Traces of the distribution of extracellular Vap33. (**B**) Left: Representative images of dendritic outgrowth in control and *vap33* wild-type rescue embryos at 15:00. Right: Quantification of the number of dendritic tips per aCC for the indicated genotypes. The statistical significance was tested to a control ① sample with a two-tailed Student’s t-test (^*^, *p* < 0.05). Data are shown as mean ± SEM. Scale bars, 5µm. (**C**) A model for aCC dendrite development.

Our studies demonstrate that Vap33 would be secreted to the extracellular space and distributed the longitudinal connectives. *vap33* can also have cell-autonomous functions, for example, in the larval neuromuscular junction (NMJ) (Pennetta et al., 2002). At the NMJ, Vap33 enables dynamic reorganization of microtubules to modulate the number of synaptic boutons.

Here, to determine the location(s) where vap33 acts to promote aCC dendritic outgrowth, we carried out rescue experiments in homozygous *vap33* null mutants. We expressed a *UAS-vap33* transgene under the control of *eve-GAL4*. We found that the aCC-specific expression of *vap33* could not rescue the mutant phenotype (**Figure S5B**). In contrast, when expressing *vap33* in all neurons except aCC (we combined the pan-neuronal *elav-GAL4* driver with the *eve-GAL80* repressor), we could rescue the phenotype (**Figure 7B** ③). These experiments suggest that *vap33* promotes dendritic outgrowth in a non-cell-autonomous manner.

Note that longitudinal axon tracts visualized by antibodies to HRP or Fasciclin II in *vap33* mutants are indistinguishable from wild-type in the CNS (**Figure S5C**). Therefore, aCC dendrite phenotypes in *vap33* could not be a secondary consequence of CNS defects; instead, *vap33* would be explicitly required for dendritic outgrowth at this developmental stage.

### *vap33* interacts genetically with *eph*

As seen in **Figure S5A**, mutants lacking *vap33* have similar phenotypes to those in *eph* mutants, suggesting that Vap33 may act as a ligand for Eph in promoting dendritic outgrowth. To determine if Vap33 and Eph function in the same signaling pathway, we examined genetic interactions between loss-of-function mutations between *vap33* and *eph*. We counted the dendrite tip number in a double heterozygous combination of the two mutants. We observed that the double mutants significantly reduce the tip number (**Figure S5D**), whereas neither heterozygous mutation alone affects dendrite development (**Figure S5D**). These data indicate that Vap33 functions together with Eph to promote dendritic outgrowth. We also assessed whether the *vap33* null mutation disrupts the pattern of Cdc42 activation. We found that the level of Cdc42 activation was significantly decreased (**Figure S5E**), as observed in *vav* or *eph* mutants. Hence the activation defects of Cdc42 in the *vap33* mutants indicate an important role of Vap33 in the proper activation of Cdc42 and raise the possibility that Vap33 regulates an aspect of Eph-dependent dendritic outgrowth.

## DISCUSSION

In this study, we investigate the molecular mechanism that underlies the outgrowth of primary dendrites. We show here that the Eph receptor plays a critical role in the process. The activation of the receptors would elevate the intrinsic kinase activity and tyrosine autophosphorylation. This phosphorylation in turn leads to the recruitment of Vav to Eph tyrosine residues via the Vav SH2 domain. The membrane localization of Vav is necessary and sufficient for dendriti outgrowth, suggesting that Vav provides a molecular link between Eph activity and Cdc42-dependent dendritic outgrowth. Analysis of *vap33* mutants has further revealed the defects in Cdc42 activation and outgrowth of primary dendrites, suggesting Vap33 may play an upstream role in Eph receptor signaling. These findings indicate that the membrane localization of Vav leads to Cdc42 activation regulated by Eph and Vap33 (**Figure 7C**).

### Eph receptor acts with Vav to regulate Cdc42 activity

Here we describe a novel function of the Rho family GEF Vav in *Drosophila*. Several observations are consistent with the notion that Vav couples the Eph receptor to the activation of Cdc42 at the onset time point of dendritic outgrowth: (1) Eph and Vav are present at 13:00 AEL in the region where Cdc42 becomes activated; (2) Vav physically associates with Eph; (3) A similar dendrite phenotype is observed when the *eph* or *vav* function is perturbed; (4) The domains in Vav and Eph that mediate their interactions are critical for the function in dendritic outgrowth; (5) Overexpression of membrane-anchored Vav is sufficient to rescue the dendritic outgrowth defects in *eph* mutants, whereas a GEF-inactive version of Vav fails to rescue the mutant phenotype; and (6) The activation of Cdc42 is disrupted in loss-of-function *vav* or *eph*. These findings suggest that Vav plays an essential role in Eph-induced Cdc42 activation, leading to an initial outgrowth of primary dendrites in the aCC motoneurons.

The small GTPases of the Rho family, Cdc42 and Rac, share 70% amino acid sequence identity. Greenberg and colleagues have previously shown that Vav-dependent Eph signaling promotes the activation of Rac for growth cone repulsion of retinal ganglion cells (RGCs) in the mouse (Cowan *et al*., 2005). In this cellular context, Rac activity is critical to promoting the internalization of the plasma membrane and contractility of the actin cytoskeleton. In contrast, our studies provide strong evidence that Vav-dependent Cdc42 activity is required to initiate dendritic outgrowth. It is conceivable that Vav activates either Cdc42 or Rac at different subcellular locations, specifying distinct downstream pathways. These differences eventually lead to the actin cytoskeletal rearrangement to specific shapes of neurons. One way to adjust the preference for Rac over Cdc42 might arise from the post-translational modification of Vav. For example, Ephexin GEF, another GEF family for Rho GTPases, can activate multiple Rho family members (RhoA, Rac1, and Cdc42) (Sahin et al., 2005; Shamah et al., 2001). When phosphorylated, however, Ephexin boosts RhoA activation relative to Cdc42 and Rac (Sahin *et al*., 2005). Phosphorylation of Vav has been reported to primarily activate Rac (Cowan *et al*., 2005). However, whether Vav phosphorylation alters the preference is currently unknown. It is tempting to speculate that, in the absence of phosphorylation, Vav may enhance activity to Cdc42 but not Rac, thus leading to dendritic outgrowth. If this is the case, Vav phosphorylation must be locally regulated. In the future, it will be critical to assess whether the kinase activity of Eph or other kinases modulates the preference of Vav GEF activity.

### Vap33 controls the outgrowth of primary dendrites

Eph receptors, associated with Eph ligands (i.e., Ephrin and Vap33), appear to mediate a wide variety of biological processes in developing brains. In the past decades, extensive studies have focused on the specification of axonal pathfinding (Bashaw and Klein, 2010; Kolodkin and Tessier-Lavigne, 2011), the establishment of topographic projection (Cang and Feldheim, 2013; Feldheim and O’Leary, 2010; Flanagan, 2006; McLaughlin and O’Leary, 2005), and dendritic spinogenesis (Hruska and Dalva, 2012; Klein, 2009; Lemke and Reber, 2005; Thompson, 2003). In contrast, we just started to form our knowledge about a specific requirement for Eph signaling in the formation of dendritic processes. For example, our recent studies have suggested a role for *Drosophila* Ephrin/Eph in olfactory map formation (Anzo *et al*., 2017). An Ephrin/Eph interaction is required to demarcate borders between dendrites of neighboring glomeruli in adult antennal lobes. In this instance, Ephrin and Eph can act as a ligand and receptor pair to mediate dendrite repulsion. Although the examinations in *Drosophila* have detected only repellent effects of Eph signaling (Anzo *et al*., 2017; Bossing and Brand, 2002; Boyle *et al*., 2006; Dearborn et al., 2002; Sardana *et al*., 2018), many studies in the developing and mature nervous system in vertebrates have shown that Eph receptors are bifunctional (i.e., having attractive and repulsive functions), regulating numerous patterning and morphological processes in the nervous system. A crucial feature of this bifunctionality, in the case of *Drosophila*, is that it might be ligand-dependent. We show that the Eph receptor can induce Vap33-dependent outgrowth of primary dendrites in the aCC motoneurons. Meanwhile, it is possible that when aCC dendrites respond to Ephrin, Ephrin/Eph signaling causes a repellent effect on dendritic outgrowth. Because these ligands are similarly present in the CNS (shown in **Figure 7A** and (Boyle *et al*., 2006)), it will be important to determine whether Ephrin and Vap33 differentially act on subcellular locations in the same neurons or different subsets of neurons.

Vap33 is composed of an MSP (major sperm protein) domain, a coiled-coil domain, and a transmembrane domain (Kamemura and Chihara, 2019). *vap33* has cell-autonomous functions in the maintenance of cellular homeostasis (Kamemura and Chihara, 2019; Lev et al., 2008; Petkovic et al., 2020), including lipid transport, microtubule reorganization, and unfolded protein response. Considerable evidence suggests that Vap33 acts as an extracellular signaling molecule as well (Han et al., 2012; Tsuda *et al*., 2008). For example, Han *et al*. recently reported that Vap33 sends signals through the Eph receptor to mitochondria in *Drosophila* muscles (Han *et al*., 2012). In this case, the neuronal Vap33 is likely cleaved, and the MSP domain is secreted into the extracellular environment. The secreted MSP functions non-cell-autonomously as a ligand that influences mitochondrial networks in muscles. Consistent with the signaling mechanism, our genetic analysis suggests the non-cell-autonomous effect of Vap33 in the aCC motoneuron. However, whether Vap33 acts directly or indirectly via the Eph receptor is not clear in the context of aCC dendritogenesis. One intriguing possibility is that an Eph-expressing aCC interacts with the secreted MSP domain of Vap33. Vap33 cleavage would be temporally regulated to yield MSP domain secretion at the onset of dendritic outgrowth. Alternatively, membrane-spanning Vap33 on the surface of neighboring cells might confer specific Eph activation to aCC. In addition, Vap33 binding to other guidance receptors (such as Robo and Lar receptors) (Han *et al*., 2012) might indirectly affect Eph signaling. Additional studies with genetic and biochemical analysis of Vap33 are required to determine how Vap33 controls the outgrowth of primary dendrites in aCC.

### Cdc42 functions in aCC to initiate dendritic outgrowth

One key question is how Cdc42 activation induces the outgrowth of primary dendrites. The control of dendritic outgrowth would result from localized modulation of the actin cytoskeleton. However, the intracellular mechanisms involved in coupling Cdc42 activation to changes in the actin cytoskeleton have remained unsolved. We previously demonstrated that at 11:00 AEL, before the outgrowth of primary dendrites, Dscam1 is concentrated at the aCC-MP1 contact site, anchoring Pak1 via Dock at a specific membrane domain in primary dendrites (Kamiyama *et al*., 2015). When Cdc42 is activated at 13:00 AEL, its interaction with Pak1 is confined to the vicinity of the localized Dscam1/Dock/Pak1 complex. Under this condition, we speculate that active Pak1 would regulate other downstream kinases (e.g., Lim-kinase and myosin-regulatory light chain) to elicit local changes in actin polymerization and to trigger spatially restricted extension of primary dendrites.

Given the critical roles of Cdc42 in regulating a wide variety of cellular processes, in regulating *de novo* actin polymerization, and now in mediating the Vap33/Eph/Vav pathway, one would expect that loss-of-function mutations of these signaling molecules lead to complete disruption of dendritic processes. For example, we have previously shown that dendritic outgrowth is disrupted entirely in *dscam1* homozygotes (Kamiyama *et al*., 2015). However, embryos homozygous for any of the *vap33, eph, vav*, and *cdc42* null alleles exhibit a mild phenotype. These findings suggest that perhaps an alternative pathway mediates dynamic remodeling of the cytoskeleton via Dscam1, thereby compensating for the loss of function of these genes. Consistent with this view, we have previously reported that Dscam1 can physically interact with the tubulin folding cofactor D (TBCD) (Okumura et al., 2015), one of the five TBCs in *Drosophila*. TBCD-dependent Dscam1 function is essential for dendrite arborization and axon maintenance of second-order olfactory projection neurons. We thus anticipate that the Dscam1-mediated dendrite growth-promoting pathway may additionally involve the regulation of microtubule dynamics (**Figure 7C**).

### Primary dendrites form through a series of choreographed signaling events

Significant progress has been made in identifying extracellular cues and cell surface receptors that regulate neuronal development. However, a key, unsolved issue is how cell surface receptors promote diverse cellular processes to create specific responses such as axon guidance, dendritic outgrowth, and synapse formation. With a limited number of surface receptors expressed, a neuron must utilize a combination of multiple receptors (i.e., guidance receptors, cell adhesion molecules, and transmembrane proteins) in determining a specific cellular process in the developing CNS. Our findings suggest that two guidance receptors transduce localized signals to modulate cytoskeletal organization spatiotemporally in the region of primary dendrites. Specifically, we find that the Dscam1 and the Eph receptor signaling pathways are both required for Cdc42-induced dendritic outgrowth but that these receptors appear to regulate different aspects of outgrowth signaling. The Dscam1 signaling pathway controls the location of dendritic outgrowth (**Figure 7C**). Before dendritic outgrowth at 11:00 AEL, an aCC growth cone expressing Dscam1 projects into the periphery. The neuron meets the MP1 neuron expressing Dscam1. Inter-neuronal interactions between Dscam1 receptors induce spatially restricted localization of Dscam1 at the contact point. This contact immobilizes Pak1 to the plasma membrane via Dock. The Eph signaling pathway, on the other hand, defines the time point of dendritic outgrowth (**Figure 7C**). At 13:00 AEL, the Vap33/Eph complex recruits Vav to the membrane, where it could then locally stimulate Cdc42. This eventually leads to Pak1 kinase activation, which is necessary for the actin assembly required for the outgrowth of primary dendrites.

We importantly note that *dscam1, dock*, and *pak1* are not necessary for Cdc42 activation because, in the previous report, we demonstrated that levels of Cdc42 activation were unaltered in these mutants (Kamiyama *et al*., 2015). Therefore, Cdc42 would be activated solely via Vav downstream of Eph independently from the Dscam1 pathway.

### Limitations of study

Our study has uncovered an important and novel function of Eph signaling in restricting dendritic outgrowth to an appropriate time. Collectively, we demonstrate that Vap33/Eph/Vav/Cdc42, in a combination of Dscam1/Dock/Pak1 located at the aCC-MP1 contact site, spatiotemporally provides the instructive forces to promote the outgrowth of primary dendrites. However, it is not clear if a combination of these guidance receptors can subserve the outgrowth of primary dendrites in different organisms or if the combination is involved in other steps in the processes of neuronal morphogenesis, such as endocytosis, local protein translation, and protein degradation in this organism. Given the versatility of these signaling pathways, these are likely to be differentially adopted in cellular contexts for specific purposes.

## Acknowledgements

We thank Edward Kipreos, Kota Banzai, Kousuke Kamemura, and members of the Kamiyama Lab for their insightful comments on the manuscript. We appreciate the gifts of reagents from Thomas Kidd, Hiroshi Tsuda, and Marianne Malartre. Stocks from the Bloomington *Drosophila* Stock Center (NIH P40OD18537) were also used in this study. This work was supported by an NIH R01 NS107558 (to Y.N., R.K., M.F., and D.K.) and UGA start-up funds (to D.K.), and JSPS KAKENHI (21H02479) and the Astellas Foundation for Research on Metabolic Disorders (to T.C.).

## Author contributions

T.C. and D.K. designed the project. D.K., Y.N., and R.K. performed experiments. R.K. and M.F. produced reagents. D.K., Y.N., and R.K. analyzed the data. D.K. wrote the manuscript. D.K., Y.N., R.K., M.F., and T.C. edited the manuscript.

## EXPERIMENTAL PROCEDURES

### Molecular cloning

Complementary DNA sequences of *vav* and *eph* (LD25754 and RE61046) (*Drosophila* Gene Collection, BDGP) were PCR amplified and subcloned into *pACUH-superfolderGFP* (*sfGFP*) to generate C-terminal *sfGFP* fusions using the NotI restriction site. The *pACUH-sfGFP* vector includes the *sfGFP* sequence in *pACUH* (Addgene) at the *Not*I and *Xba*I sites. To create deletion constructs (*vav*^*ΔSH2*^ and *eph*^*Δkinase*^), we performed site-directed mutagenesis using the Q5 Site-Directed Mutagenesis Kit (New England Biolabs). We deleted a DNA sequence encoding the SH2 domain of Vav (amino acids 622-706) or the kinase domain of Eph (amino acids 692-950) from *pACUH-vav-sfGFP* or *pACUH-eph-sfGFP*, respectively. To replace tyrosine (Y) with phenylalanine (F), we designed two pairs of mutagenesis primers. We performed two rounds of mutation reactions and introduced four Y-to-F substitutions to the juxtamembrane region of Eph. A synthesized DNA fragment (Integrated DNA Technologies) containing either *myr-vav*^*GEF-PH*^ or *myr-vav*^*GEF[AA]-PH*^ was cloned into the *Eco*RI and *Not*I sites of *pACUH-sfGFP*. Those Vav^GEF-PH^ constructs comprise a GEF-PH domain only (amino acids 220-541).

For co-immunoprecipitation assays, we designed plasmids to produce FLAG-tagged or Fc-tagged proteins. *vav-3xFLAG* and *vav*^*ΔSH2*^*-3xFLAG* fragments were PCR amplified from *pACUH-vav-sfGFP* and *pACUH-vav*^*ΔSH2*^*-sfGFP*. Then, we introduced each fragment to the *pACUH* vector through the *Not*I and *Xba*I sites, constructing *pACUH-vav-3xFLAG* and *pACUH-vav*^*ΔSH2*^ *-3xFLAG*. Human *Fc* fragments were PCR amplified from the *pIB-Fc* vector (a gift from Thomas Kidd). *eph-IC* and *eph*^*Y-F*^*-IC* were generated via PCR from the full-length *eph* and *eph*^*Y-F*^. *eph-IC-Fc* and *eph*^*Y-F*^*-IC-Fc* were assembled into *Not*I- and *Xba*I-digested *pACUH. eph-IC* encodes a fragment of the C-terminal region (amino acids 629-1047).

Standard subcloning, PCR amplification (PrimeSTAR Max polymerase, Takara Bio USA), and In-Fusion cloning (In-Fusion Snap Assembly Master Mix, Takara Bio USA) were used to generate all plasmids and intermediates. The oligonucleotides and gene fragments used are listed in **Table S2**.

### Recombineering of a *GFP-vav* fusion

A *GFP-vav* fusion construct, *P[acman-GFP-vav]*, was generated using a recombineering strategy from a *P[acman] BAC* clone, *Ch322 143P01* (The BRB Preclinical Repository). Recombineering was performed according to Venken *et al*. (Venken et al., 2006). Briefly, a *GFP-kanamycin* marker was PCR amplified from a template vector (*PL452*, Addgene). The primers include homology arms consisting of 50 nucleotides upstream and downstream from the start codon of the *vav* gene. The set of primers enables N-terminal tagging of the *vav* gene.

Next, the PCR product was transformed into competent bacteria, SW106 (the BRB Preclinical Repository), that harbor *P[acman-vav]*. Recombination between the homology arms and *vav* was then selected using a kanamycin medium. Subsequently, we induced Cre recombinase expression in the medium supplemented with L-arabinose and deleted the kanamycin-resistant cassette from the BAC plasmid. After plasmid extraction, the *GFP-vav* insertion site was verified by sequencing across the 5’ and 3’ homology arm junctions.

### Transgenic flies

After being sequenced, all plasmids were injected into *attP* (*attp2, attp40*, or *VK00027*) embryos expressing *ΦC31* by Rainbow Transgenic Flies, Inc.

### Fly stock

Standard *Drosophila* genetic techniques were used for all crosses. All flies were raised at 25°C. The following lines were obtained from the Bloomington *Drosophila* Stock Center: *eve-GAL4* (BDSC stock no. 7470 and 7473), *elav-GAL4* (8760), *UAS-sif RNAi* (61934), *UAS-ziz RNAi* (54817), *UAS-rhogef3 RNAi* (42526), *UAS-rtgef RNAi* (32947), *UAS-rhogef4 RNAi* (42550), *UAS-pbl RNAi* (36841), *UAS-cdep RNAi* (31168), *UAS-psgef RNAi* (44061), *UAS-gefmeso RNAi* (42545), *UAS-trio RNAi* (43549), *UAS-unc-89 RNAi* (34000), *UAS-rhogef64c RNAi* (31130), *UAS-pura RNAi* (58270), *UAS-zir RNAi* (53946), *UAS-spg RNAi* (35396), *UAS-exn RNAi* (33373),*UAS-vav RNAi* (39059), *UAS-rhogef2 RNAi* (34643), *UAS-eph RNAi* (60006), *UAS-vap33 RNAi* (77440), and *UAS-FLAG-vap33-HA* (39682). *vav*^*2*^ and *vav*^*3*^ mutants were provided by Dr. Marianne Malartre, The French National Centre for Scientific Research, France. The genotypes used in each experiment are listed in **Table S1**.

### Immunohistochemistry

Dissected embryos were fixed with 4% paraformaldehyde in phosphate buffered saline (PBS) for 10 min, washed in TBS (0.1% Triton X-100 in PBS), blocked in 10% bovine serum in TBS for 1hr, and finally incubated with a primary antibody overnight at 4°C. The following primary antibodies were used: mouse anti-myc (Developmental study Hybridoma Bank, 1:100), chicken anti-GFP (abcam, 1:500), Alexa647-conjugated goat anti-HRP (Jackson ImmunoResearch Labs, 1:500). We then washed and incubated embryos in a secondary antibody conjugated to a fluorescent dye (Thermo Fisher Scientific, 1:500) for 2hr at room temperature. After extensive washes with TBS, embryos were mounted in 50% glycerol for slide preparation and imaged using a confocal microscope. At least ten different embryos per experimental condition were imaged to quantify fluorescence signals. For extracellular Vap33 localization studies, we omitted permeabilization before antibody staining. Similar immunostaining was performed as described previously (Tsuda *et al*., 2008). This way, we can specifically label extracellular Vap33 to distinguish them from the intracellular pool. Rabbit anti-Vap33 antibodies were used at 1:10000 (a gift from Hiroshi Tsuda)

### Lipophilic dye labeling

*In vivo* labeling of the aCC motoneurons with a lipophilic dye was performed as previously described (Inal *et al*., 2020).

### Fluorescence Imaging

Fluorescence images were taken using an inverted microscope (Ti-E, Nikon) with a 100× 1.45 NA oil immersion objective (Plan Apo, Nikon). The microscope was connected to a Dragonfly Spinning disk confocal unit (CR-DFLY-501, Oxford Instruments). Three excitation lasers (40mW 488nm, 50mW 561nm, and 110mW 642nm lasers) were coupled to a multimode fiber passing through an Andor Borealis unit. A dichroic mirror (Dragonfly laser dichroic for 405-488-561-640) and three bandpass filters (525/50nm, 600/50nm, and 725/40nm bandpass emission wheel filters) were placed in its imaging path. Images were recorded using an electron multiplying charge-coupled device camera (Andor iXon, Oxford Instruments) controlled with Fusion software (Oxford Instruments).

### Cell culture

*Drosophila* S2R+ cells were maintained in an SFX-INSECT culture medium (Hyclone) at 25°C. Cells were transfected with 300ng of *pACUH* constructs using 2.5µl of Effectene (Qiagen). Transiently transfected cells were induced to express by co-transfecting 300ng of *pAC-GAL4* (Addgene) and analyzed at 48hr post-transfection.

### Co-IP assay

S2 cells expressing *pACUH* constructs were lysed in 450µl of 1% NP40 lysis buffer. Lysates were incubated with 5µl of Pierce Protein A/G Magnetic Beads (Thermo Fisher Scientific) overnight at 4°C with continuous rotation. The samples were washed three times with PBS and eluted with an SDS loading buffer. Samples in the buffer were subjected to SDS-PAGE. The proteins were transferred onto PVDF membranes (MilliporeSigma). The membranes were blocked at room temperature for 1hr in 4% skimmed milk in TBS (0.1% tween in PBS) and probed with 1:1000 diluted mouse anti-FLAG (Sigma-Aldrich). Anti-mouse secondary antibodies conjugated to horseradish peroxide (Thermo Fisher Scientific) were used at a dilution of 1:10000 for 1hr at room temperature. Blots were developed with a chemiluminescent substrate (SYNGENE) and imaged using the ChemiDoc Imaging System (BIO-RAD).

### FRET measurement

FRET was measured using an upright microscope (Axio imager Z2; Carl Zeiss) with a 63× 1.4 NA oil immersion objective (Plan Apo, Carl Zeiss). The upright microscope was attached to an LSM 880 scanning head (Carl Zeiss) with a 32-channel GaAsP spectral photomultiplier tube detector. We used the 458nm line of an argon laser with the main beam filleter MBS-458/514. Emitted light was detected using two emission filters (454-518nm barrier filter for CFP and 517-605nm barrier filter for YFP). Images were captured using Zen software (Carl Zeiss). Raw images were corrected by subtracting the background fluorescence intensity of non-labeled cells. Yellow/cyan emission ratios were calculated along the length of an individual axon.

### Quantification of fluorescence signals

Optical sections of 0.5µm were taken across the depth of the embryos and were reassembled with ImageJ software (The National Institutes of Health). Identical laser power, gain, and exposure time settings were used to collect data. The final pictures were shown as average intensity projections. When quantifying Vap33, Eph, and Vav signal levels in the CNS, 30µm-long lines were drawn perpendicular to the midline of an image of a z projection. Pixel intensity along this line was average for 20µm long along the anterior-posterior axis. We plotted these values in line charts according to distance. To quantify fluorescence signals in the aCC motoneurons, a high-resolution image of each neuron was background-subtracted and threshold-segmented. Then, the mean fluorescence signals within each neuron were measured. Per genotype, >10 neurons were quantified and averaged. Again, these average values were plotted at 5 to 30 µm from the midline in line charts.

### Statistical analysis

Statistical analysis was performed using OriginPro (OriginLab) and Excel (Microsoft). Data are presented as means ± standard errors of the means (SEMs) in all figures. To perform statistical analysis on the amount of protein at the site of aCC dendritogenesis, we quantified it by fitting each line chart to a Gaussian function and using the resultant area. The uncertainty of this measurement was estimated by bootstrapping individual charts. The type of statistical test performed and replicate number for each experiment are listed for each analysis. Significance was classified as *p* < 0.05.

