## Supplementary Information for "The Vap33/Eph/Vav/Cdc42 complex confers temporal specification to the outgrowth of primary dendrites in Drosophila neurons"

**Figure S1:**

*Drosophila vav* is a Cdc42 GEF involved in dendritic outgrowth in aCC

**Figure S2:**

Vav distribution changes with different developmental stages

**Figure S3:**

Alignment of the juxtamembrane sequences from *Drosophila* Eph and mouse EphA4

**Figure S4:**

*eph* functions in the same genetic pathway as *vav*

**Figure S5:**

Vap33 activates Cdc42, which promotes dendritic outgrowth

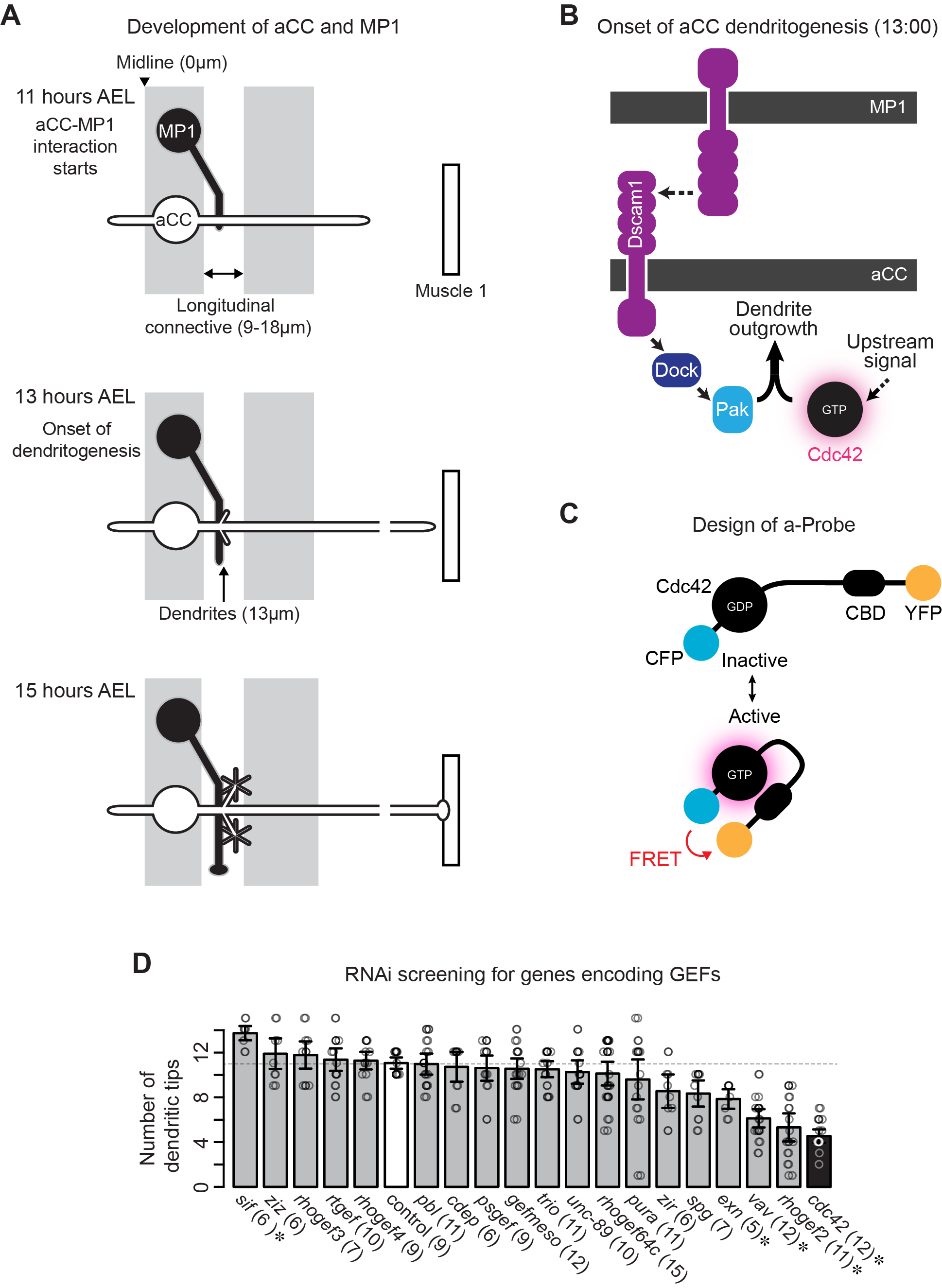

**Figure S1, related to Figure 1 and 2.­­­­­­ *Drosophila vav* is a Cdc42 GEF involved in dendritic outgrowth in aCC**

(**A**) Schematic showing development at 11:00, 13:00, and 15:00 AEL of the aCC and MP1 neurons in the embryonic CNS. (**B**) The schematic represents our understanding of how the Dscam1/Dock/Pak1 complex promotes aCC dendritogenesis at the aCC-MP1 contact site at 13:00. (**C**) Domain composition of a Cdc42 activation probe (also called a-Probe). aProbe contains two fluorescent proteins (ECFP and Venus) on the N- and C- termini, respectively, with the Cdc42 binding domain (CBD) of Pak, Cdc42, and a flexible linker. (**D**) Results from RNAi screens for genes encoding Cdc42 GEFs. The number of dendritic tips was quantified in individual neurons expressing *UAS-RNAi* constructs driven by *eve-GAL4*. A two-tailed Student’s t-test was used to a control sample for statistical analysis (^*^, *p* < 0.05).

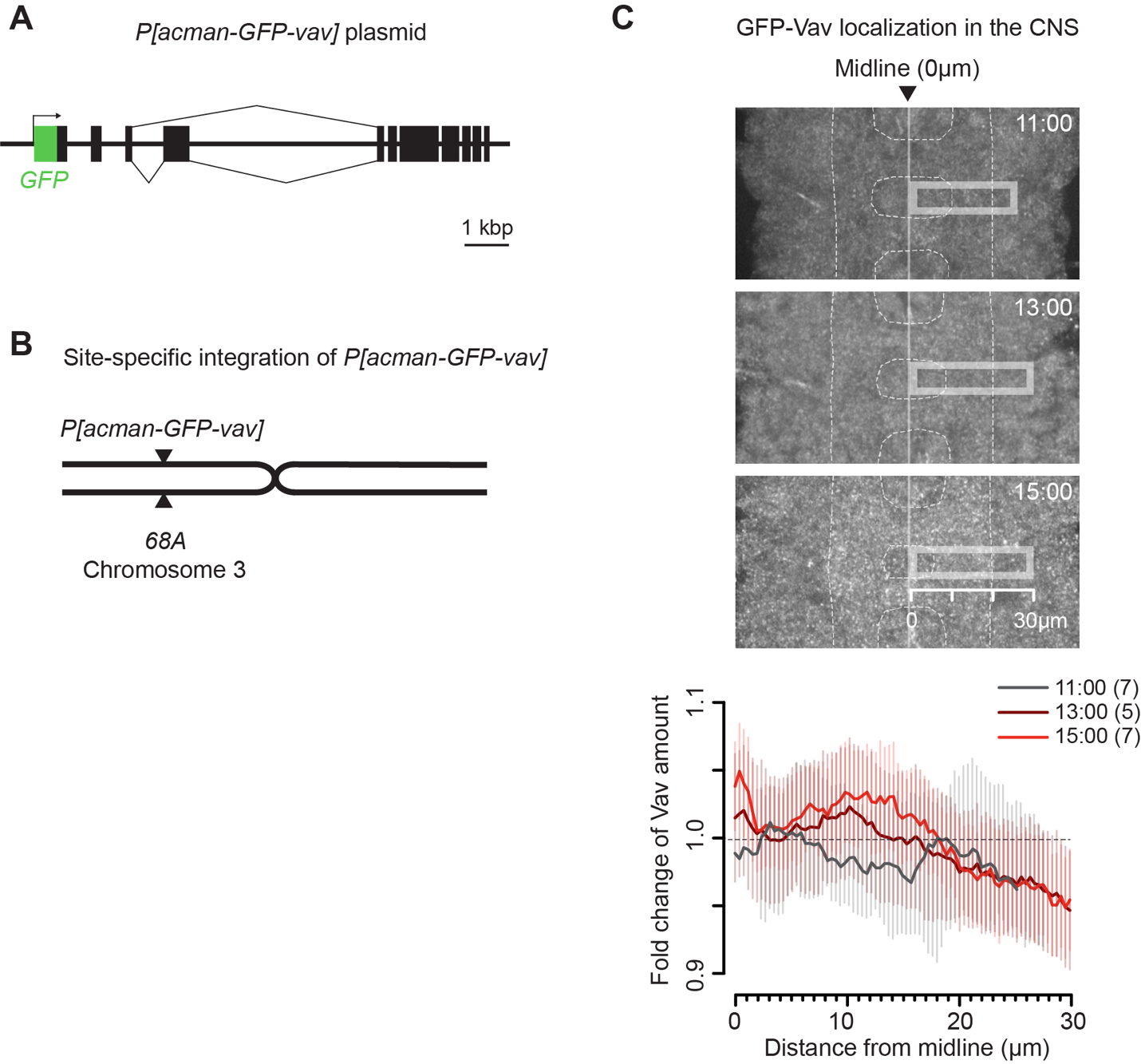

**Figure S2, related to Figure 3. Vav distribution changes with different developmental stages**

(**A**) Schematic of the genomic structure of the *vav* gene in a BAC plasmid. A *P[acman-GFP-vav]* construct was made by inserting *GFP* immediately after the start codon of *vav* via recombineering. (**B**) The construct was inserted into the cytological location 68A of the third chromosome using site-specific integration. (**C**) Top: Fluorescence images of anti-GFP staining of *P[acman-GFP-vav]* embryos at the indicated developmental stages. Bottom: Traces of GFP-Vav distribution in the CNS.

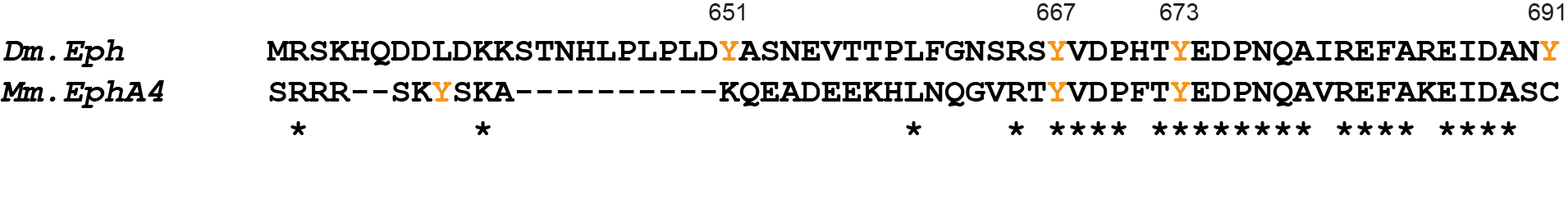

**Figure S3, related to Figure 4. Alignment of the juxtamembrane sequences from *Drosophila* Eph and mouse EphA4**

The juxtamembrane segment sequences from *Drosophila* Eph and mouse EphA4 are aligned using ClustalW2. Tyrosine residues are highlighted in orange. An asterisk denotes a conserved residue.

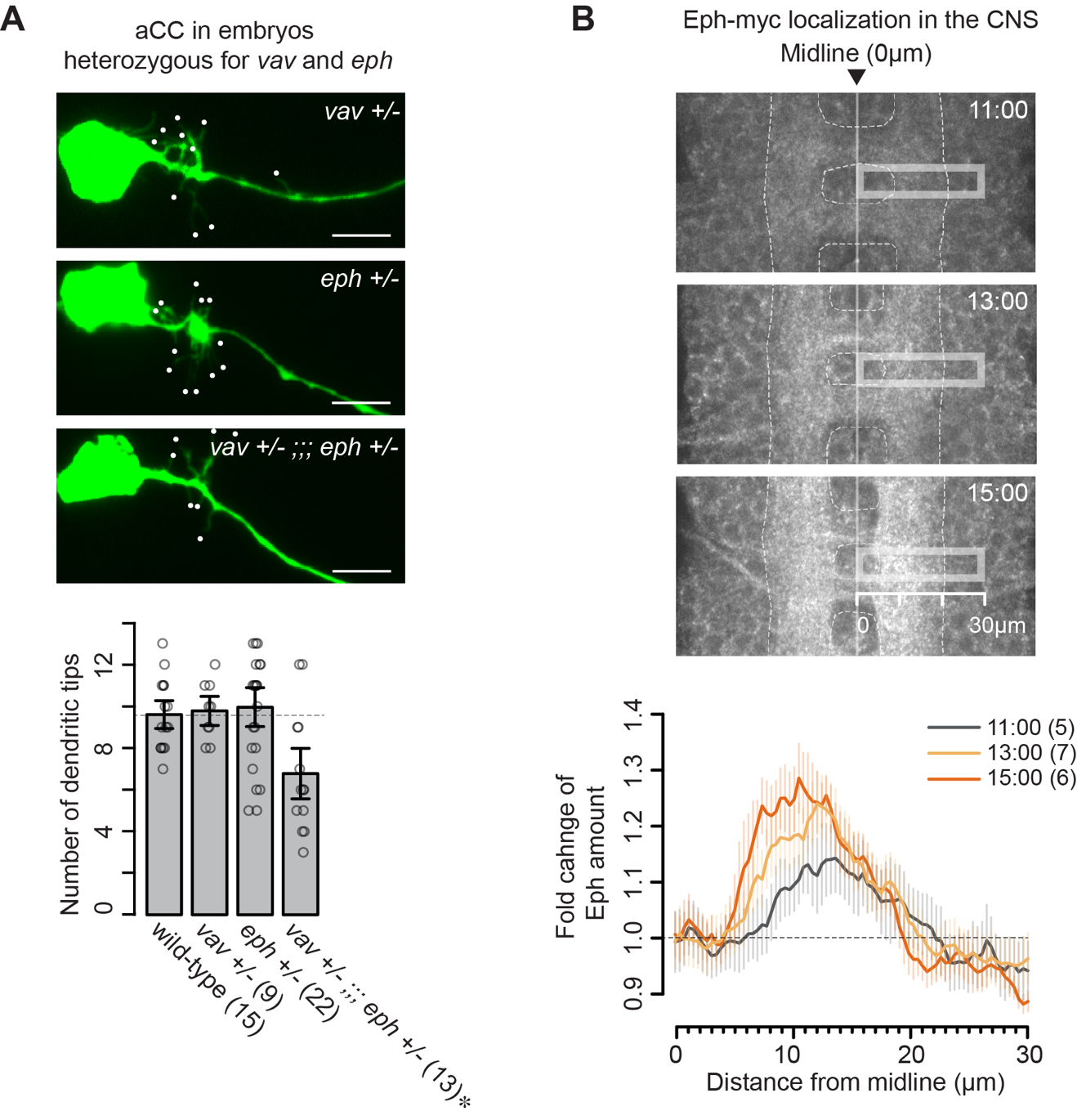

**Figure S4, related to Figure 5. *eph* functions in the same genetic pathway as *vav***

(**A**) Top: Representative images of dendritic outgrowth in embryos heterozygous for *vav*, heterozygous for *eph*, and doubly heterozygous for *vav* and *eph* at 15:00. Bottom: Quantification of the number of dendritic tips per aCC for the indicated genotypes. The statistical significance was tested to a control sample (wild-type) with a two-tailed Student’s t-test (^*^, *p* < 0.05). (**B**) Top: Fluorescence images of anti-myc staining of *Eph-myc* embryos in the indicated developmental stages. Bottom: Traces of Eph-myc distribution in the CNS. Scale bars, 5μm.

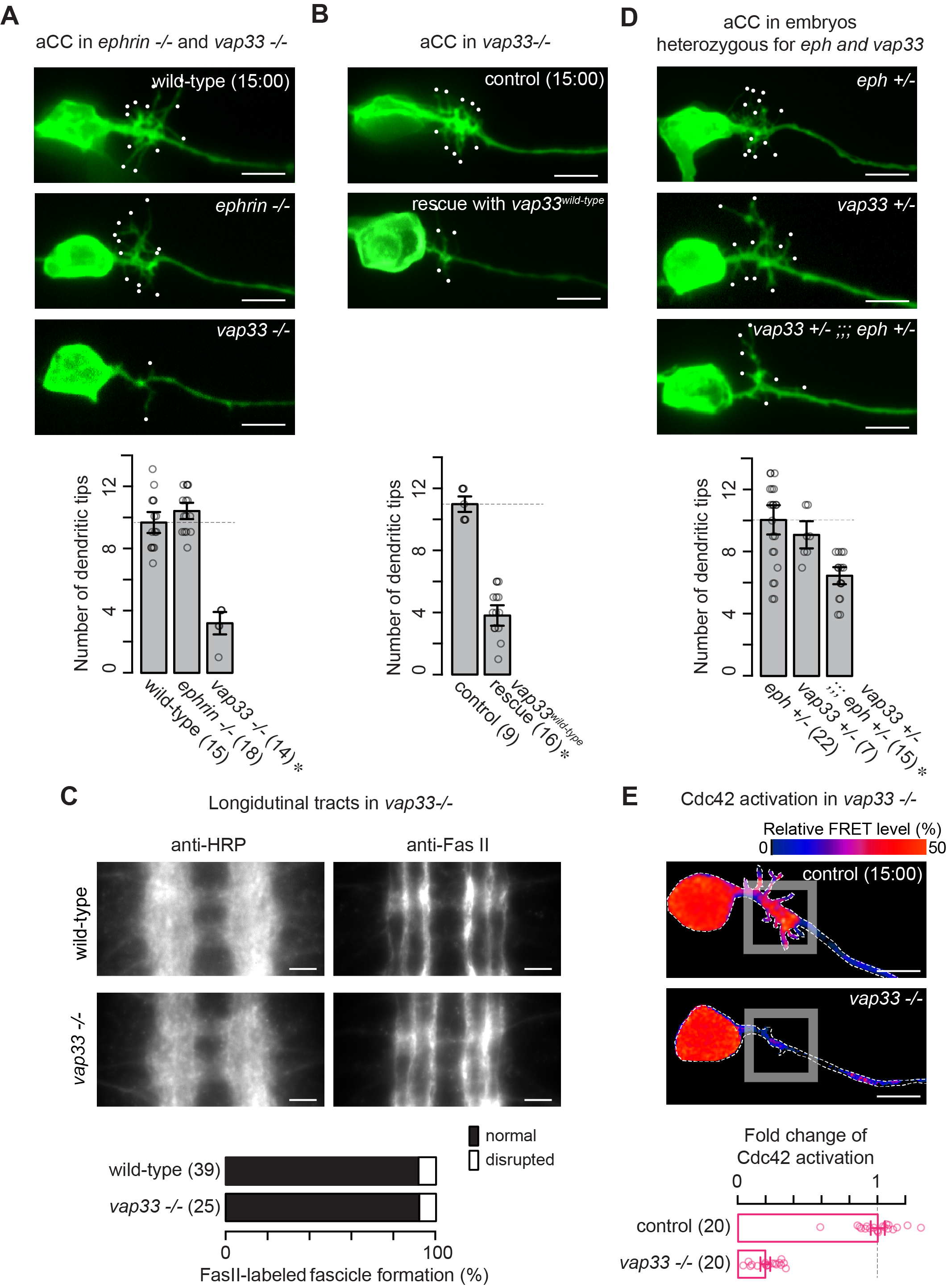

**Figure S5, related to Figure 7. Vap33 activates Cdc42, which promotes dendritic outgrowth**

(**A** - **B**) Top: Representative images of dendritic outgrowth in *ephrin* null, *vap33* null, and *vap33* wild-type rescue embryos at 15:00. (**D**) Top: Representative images of dendritic outgrowth in embryos heterozygous for *eph*, heterozygous for *vap33*, and doubly heterozygous for *eph* and *vap33*. (**A** - **B** and **D**) Bottom: Quantification of the number of dendritic tips per aCC for the indicated genotypes. A two-tailed Student’s t-test was used to a control sample for statistical analysis (^*^, *p* < 0.05). (**C**) Top: The CNS of wild-type or *vap33* null embryos stained with anti-HRP or Fas II at 15:00. Bottom: Quantification of axon growth and guidance defects in wild-type or *vap33* null embryos. No prominent disruption in the longitudinal tracts was detectable. (**E**) Top: Pseudocolor images of aProbe responses in control and *vap33* null embryos. Bottom: Quantification of the level of Cdc42 activation per aCC for the indicated genotypes. Scale bars, 5μm in (**A**, **B**, **D** and **E**) and 10μm (**C**)­­­.
